# Generating complex patterns of gene expression without regulatory circuits

**DOI:** 10.1101/2020.11.25.398248

**Authors:** Sahil B. Shah, Alexis M. Hill, Claus O. Wilke, Adam J. Hockenberry

## Abstract

Synthetic biology has successfully advanced our ability to design and implement complex, time-varying genetic circuits to control the expression of recombinant proteins. However, these circuits typically require the production of regulatory genes whose only purpose is to coordinate expression of other genes. When designing very small genetic constructs, such as viral genomes, we may want to avoid introducing such auxiliary gene products while nevertheless encoding complex expression dynamics. To this end, here we demonstrate that varying only the placement and strengths of promoters, terminators, and RNase cleavage sites in a computational model of a bacteriophage genome is sufficient to achieve solutions to a variety of basic gene expression patterns. We discover these genetic solutions by computationally evolving genomes to reproduce desired gene expression time-course data. Our approach shows that non-trivial patterns can be evolved, including complex patterns where the relative ordering of genes by abundance changes over time. We find that some patterns are easier to evolve than others, and comparable expression patterns can be achieved via different genetic architectures. Our work opens up a novel avenue to genome engineering via fine-tuning the balance of gene expression and gene degradation rates.

**Author summary:** Viruses that infect bacteria, commonly called bacteriophages, typically have small genomes that encode as few as 10 genes. From the perspective of understanding genome design and regulation, these organisms are important model systems. Similar to cellular species, the genes encoded on a phage genome often must be expressed at different levels and at particular times during the phage lifecycle. Given their unique size constraints, it may be advantageous for phages to accomplish differential gene expression without having to produce a variety of dedicated regulatory molecules—which are frequently encoded on the larger and more complex genomes of free-living species. Here, we use a computational simulation of phage infection coupled with an evolutionary selection algorithm to illustrate that phage genomes can encode complex time-dependent gene expression patterns without the need for dedicated regulatory molecules. We anticipate that this simulation framework may additionally aid future phage genome design and engineering efforts.

## Introduction

All genomes encode a variety of distinct RNA and protein sequences, and the production of these biomolecules is essential for the growth and survival of organisms. Critically, these various gene products are often required in different amounts relative to one-another and this demand further varies over time [1–3]. Thus, organisms must have the ability to modulate gene expression levels to meet lifestyle needs and adapt to changing environmental conditions [4–6]. Many gene-regulatory elements are well characterized and cells have been directly engineered to produce individual protein products for decades; the strength of promoters, terminators, ribosome binding sites, *etc.* can all vary over several orders of magnitude to produce different amounts of recombinant gene products [7–9].

More recently, the field of synthetic biology has developed and achieved success in engineering large, time-varying genetic circuits that are characterized by complex interactions and often require producing gene products whose sole job is to regulate the expression of other genes [10–15]. While the scale and capabilities of these applications are continually expanding [16–18], the goal of designing entire genomes from scratch presents tremendous challenges. One particular source of challenges is genome complexity—even the smallest free living organisms encode hundreds of gene products that interact with one-another and alter gene expression patterns in ways that are difficult to predict [19].

Even smaller model systems are viruses and bacteriophages. Bacteriophages are useful from an engineering and genome-design standpoint because of their comparatively small size, genetic tractability, and potential uses in medical and biotechnological applications [20–25]. Despite the limited set of genes encoded by their genomes, phages are nevertheless capable of producing complex, time-varying patterns of gene expression [26]. While some of these dynamics are governed by networks of sequence-specific promoters and transcription factors, some phages—such as the *Escherichia coli* phage T7—appear to regulate a portion of their transcriptional demand by tuning production and degradation rates [27, 28]. The full range of expression dynamics that can be achieved by varying only these mechanisms is currently unknown.

Here, we use an *in silico* model of phage infection to explore the range of gene expression complexity that can be achieved using only basic regulatory mechanisms, including transcription, transcript termination, transcript cleavage, and transcript degradation. Our approach relies on computationally evolving genomes with variable-strength promoters, terminators, and RNase cleavage sites to reproduce predefined gene expression time-course patterns. We show that this molecular-level simulation framework can discover solutions to numerous non-trivial patterns and that successfully evolved genomes for particular patterns display a variety of distinct genome architectures. Taken together, these findings lay the groundwork for future efforts to use *in silico* evolutionary design methods to engineer novel phage genomes.

## Materials and Methods

### Gene expression simulations

We used the gene expression simulation platform Pinetree [29], version 0.3.0, to simulate expression of mRNA transcripts encoded by a bacteriophage genome containing a small number of genes (between three and ten in our simulations). Our gene simulations included mRNA translation and protein production—with equal ribosome binding site strengths assigned to all genes—but for the purposes of this study we analyzed only mRNA transcript levels. Gene expression in Pinetree is stochastic, and it is modeled at the level of individual molecules. In particular, RNA polymerases are tracked individually as they attach to the bacteriophage genome and commence transcription one nucleoside at a time. Additionally, the polymerases can: i) collide with other polymerases transcribing the same genomic region, ii) fall of prematurely, iii) terminate transcription upon encountering a terminator element, or iv) read through a terminator.

In addition, RNase molecules can attach to RNAse cleavage sites within transcripts and cleave them—leading to the subsequent directional degradation of RNA fragments from the 5’ to the 3’ end. Degradation of transcripts from cleavage sites is 1000-fold more efficient than degradation of nascent transcripts from their 5’ end. A detailed explanation and justification for the directional transcript degradation model is provided in Ref. [28]. We additionally extended this prior work to allow for site-specific RNase cleavage strengths, and included this feature in the Pinetree 0.3.0 release. RNase cleavage strengths can be thought of as composite parameters that incorporate both the binding specificity of the RNase molecule and the enyzmatic cleavage rate. Either of these biochemical processes may exhibit sequence specificity, and here we did not model them individually.

Pinetree takes an arbitrary genome as input (which we note is not a sequence file but rather an input file defining the location of individual genes, promoters, terminators, etc.), as well as parameters describing rate constants and cellular-level properties of the infected host cell, such as cell volume or number of specific molecules within the cell. We kept most values at their defaults, but changed the footprint sizes of polymerase and ribosome molecules to better reflect their realistic size (35 and 30, respectively) and additionally set the polymerase copy number to a value of 4 to shorten simulation run-times. In all cases, we simulated the initial five minutes of real time infection that begins with the bacteriophage genome being injected into the infected cell, akin to the transient expression pattern displayed by a typical lytic bacteriophage.

The primary phage model we studied here consists of genomes containing three protein-coding genes, each 150 nucleotides long. Thus, each encoded protein was 50 amino acids long, and none of the proteins had any function within the simulation; we merely used them to study arbitrary gene expression regulation. We also considered genomes containing ten protein-coding genes, to investigate the scalability of our approach to larger systems.

### Evolving genomes with specific gene expression patterns

We used our stochastic model of gene expression to evolve phage genomes to display specific temporal patterns of expression. In all cases, we first defined a target phenotype as a gene expression time-course for all genes included in the genome. We then took a starting genome with a single, moderate-strength promoter upstream of the first coding sequence in the genome, and we subjected this genome to subsequent rounds of mutation and selection. Our evolutionary simulations ran for 5000 generations each, and we implemented a simulated annealing procedure that steadily increased the strength of selection with increasing number of generations. The goal with this setup was not to simulate realistic evolutionary processes of natural populations but rather to permit efficient exploration of a potentially rugged fitness landscape and thus maximize our chance of finding suitable genome architectures.

We considered three different types of mutations: addition of a regulatory element (in this case, either a promoter, terminator, or RNase cleavage site), removal of a single existing regulatory element, or modification of the binding constant for an existing regulatory element. The addition and removal functions act as insertion and deletion mutations, respectively, whereas modifications to binding strengths are analogous to point mutations that alter specific rate parameters. All protein-coding sequences are defined in the starting genome and are not altered through any mutational step.

To determine the effective ranges of individual regulatory elements as well as their starting values when they are inserted into a genome, we simulated simplified genomes and measured the expression response of downstream genes (S1 Fig) using a model containing only three coding sequences. From this qualitative analysis we chose to insert promoters at a strength of 10e6 (with minimum and maximum values set at 10e5 and 10e13, respectively), RNAse sites at 5e-3 (with minimum and maximum values set at 0 and 1), and terminators at a strength of 0.2 (with minimum and maximum values set at 0 and 1). After inserting an element, the element strength can be modified in subsequent mutational steps. Each modification is proposed by drawing a value from a normal distribution with mean of 1 and standard deviation of 0.1, and multiplying this value by the current element strength. If the proposed mutation results in an out-of-bounds value, the value is simply discarded and the process repeated until a valid proposed mutation is drawn.

Our mutational process is not meant to mimic natural evolution. Rather, *some* mutation is always proposed at every generation—either element addition, removal, or strength modification. This is accomplished by first enumerating all possible mutations given the current genome status (deletion and modification can happen to every existing element, and every non-existing element can be added) and then randomly selecting one of the possibilities from this set (*i.e.* addition of a promoter between genes 2 and 3). Only one of each element-type can exist in a particular region between two genes, so once there is a terminator between genes 2 and 3, for instance, another terminator can not be proposed in this specific location.

While some mutation will always be proposed at each generation, many of these mutations will not be accepted. We calculated the fitness of a genome from the distance between its gene expression pattern and the target pattern. Because gene expression patterns are obtained from stochastic simulations, we performed ten replicate simulations for each genome and averaged mRNA abundances across replicates before proceeding with the fitness calculations and determination of acceptance probabilities. We defined fitness in terms of the normalized root mean square error (normalized RMSE). We obtained normalized RMSE by first calculating the RMSE for a single gene *k*,

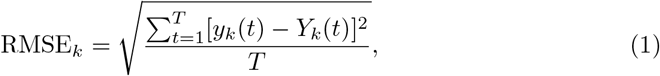

where *y*_*k*_(*t*) is the observed gene expression level of gene *k* at time *t*, *Y*_*k*_(*t*) is the target gene expression level of gene *k* at time *t*, and *T* is the total number of time steps simulated for the gene expression time course. Subsequently, we normalized each RMSE_*k*_ by the mean target abundance, 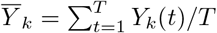, and then averaged these values across all genes,

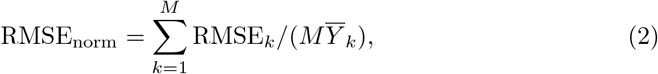

where *M* is the total number of genes in the genome. Additionally, and after analysis of several distinct patterns and simulations, we determined that normalized-RMSE values ≤ 0.1 aligned with our expectation about the suitability of individual solutions. We settled on this value and applied this threshold throughout the results whenever we refer to simulations that produce a final pattern that successfully matched the target.

To convert RMSE_norm_ into fitness *f*, we employed the Fermi function

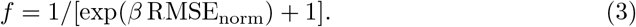

We have *f* = 1/2 when RMSE_norm_ = 0 (*i.e.* when the genome exactly displays the target expression pattern) and *f* monotonously declines to zero as RMSE_norm_ increases.

The constant *β* determines how quickly *f* declines with increasing RMSE_norm_, and thus we can vary *β* to modify the strength of selection in the simulations.

We evolved genomes using an accelerated origin–fixation model [30]. In this model, at any point in time the evolving population is represented by a single resident genotype. Whenever a novel, mutated genotypes arises, it can either go to fixation, thus becoming the next resident genotype, or it can be lost to genetic drift. Whether a new genotype is accepted or not is determined probabilistically according to the value of *P*_accept_, defined as

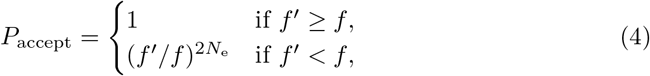

where *f* is the fitness of the resident genotype, *f′* is the fitness of the new mutant, and *N*_e_ is the simulated effective population size. We note that beneficial mutations are unconditionally accepted. This choice allows for rapid exploration of the fitness landscape while resulting in steady-state sampling identical to the one obtained when using the traditional Kimura formula for the probability of fixation [30].

To allow for even more rapid sampling of the fitness landscape, we implemented a simulated annealing procedure where selection was weak initially but became increasingly stronger towards the end of the evolution. We modified the strength of selection by varying the parameter *β* in Eq. (3), while keeping *N*_e_ constant at *N*_e_ = 1000 throughout. We varied *β* according to the following schedule: For the first 500 generations, *β* = 10^−3^, to noisily explore the fitness landscape (S2 Fig). From generation 501 to 4500, we linearly increased *β* from 10^−3^ to 1.1. Finally, from generation 4501 to 5000, we linearly increased *β* from 1.1 to 1.3. This simulated annealing approach ensured that evolutionary simulations explored a wide range of distinct genome architectures yet tended to settle on one genome architecture towards the end of each evolution. For each pattern that we explored, we performed 50 replicate simulations using the above approach.

### Accounting for possible gene re-arrangements

In our gene expression model, gene order affects observed gene expression levels, because transcription proceeds from the 5’ end to the 3’ end but can stochastically terminate at any point. Further, the genome is injected beginning at the 5’ end, which allows for polymerase binding in this region to occur before it is possible to bind promoters that are located closer towards the 3’ end of the genome. Thus upstream genes are transcribed somewhat earlier and at somewhat higher levels than downstream genes, all else being equal. Therefore, gene identity matters when comparing a genome’s expression pattern to a target pattern and it is a very different requirement to state that the first gene in a genome has to be the most expressed versus the last gene.

In practical applications of genome engineering, however, we would not normally be concerned about the order of genes in the genome, as long as each gene is expressed at appropriate levels over time. Thus, we explored *general patterns* by simulating all possible gene arrangements for each pattern. For a three-gene genome, there are six possible combinations for matching the genes in the genome to the genes described in the target expression pattern. We simulated all six variations for each pattern, and then we selected the gene arrangement that had the lowest normalized-RMSE (averaged across 50 independent simulation replicates) as the representative for that pattern. For the 10 gene simulation, we simulated only a single predetermined gene order and note that attempting to simulate all possible gene order variants would be impractical for this case.

### Removing elements with negligible effect

When analyzing final evolved genome architectures, we wanted to be certain that each regulatory element (promoter, terminator, RNase cleavage site) contained within that genome had a substantial effect on the overall expression pattern. Since rate constants for elements were allowed to evolve, it was possible that individual elements evolved towards rate constants that were so low that the elements had essentially no function. Thus, at the end of evolutionary simulations, we evaluated each element’s effect on the final gene expression pattern and removed all elements with a negligible effect.

To do so, we removed each element from the genome and then analyzed how the resulting gene expression pattern changed relative to the complete genome as it evolved. If the normalized-RMSE value remained below the threshold that we used as a quantitative indicator of a successful simulation (using a threshold of 0.1, as previously noted), we recorded the element as a candidate for removal. After evaluating all regulatory elements in this manner, we removed the element whose removal had the least impact. We then repeated this process, greedily removing elements one by one until no further elements could be removed without pushing the normalized-RMSE value above our threshold of success.

### Assessing the diversity of evolved genomes

Our evolutionary simulations generally resulted in multiple, independently evolved genomes that had different genome architectures for each target pattern. To quantify the diversity of evolved architectures, we calculated the entropy of the set of evolved solutions that were deemed successful for the given target. In this calculation, we only considered the presence or absence of individual regulatory elements (after removing insubstantial elements, as described above) in the genome while ignoring the rate constants/element strengths. We identified the unique genome architectures among the *n* successful solutions (ignoring simulations that produced a normalized-RMSE >0.1) and then determined the count *n*_*i*_ for each unique architecture *i* (so that ∑_*i*_*n*_*i*_ = *n*). We then calculated entropy *H* as

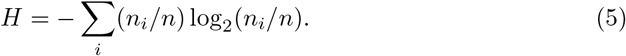

Because the logarithm is taken to base 2, the resulting entropy value is measured in units of bits.

### Data and code availability

All code and data required to reproduce this work are available at: https://github.com/SahilBShah/pinetree-evolution and also archived on Zenodo at https://doi.org/10.5281/zenodo.4592577.

## Results

### Evolutionary simulation of phage gene expression

Our goal is to understand the range of possible gene expression patterns that a phage could produce by varying only a small set of regulatory components. We reasoned that rational genome design would be difficult for all but the most trivial cases, and that fully enumerating all possible genomes is impractical. Instead, we focused on developing an evolutionary strategy to engineer *in silico* genomes to match a diverse set of target pre-defined gene expression time-courses.

This computational strategy that we developed relies on a stochastic gene expression platform (Pinetree), which simulates the process of phage infection in molecular-level detail and produces time-course data of RNA abundance as an output [29]. We built upon this software to enable evolutionary simulations, where discrete generations consist of individual simulations of phage infection and the resulting RNA species time-course is our ultimate phenotype of interest. Pinetree expects a genome as input and under our evolutionary approach this genome varies from generation-to-generation as the location and strengths of promoters, terminators, and RNase cleavage sites are subject to mutation, selection, and drift.

As a proof-of-principle, we first used Pinetree to simulate gene expression from an arbitrary example genome architecture. The positive control genome architecture and resulting gene expression pattern produced from a single simulation are shown in Fig. 1A (first panel). Using this gene expression output as our target data, we next attempted to evolve a genome from scratch that was capable of producing similar levels of gene expression over time. Our evolutionary simulation starts with a three-gene genome containing only a single promoter, whose time-course of gene expression initially bears little resemblance to the target. As generations progress, single mutations are proposed and conditionally accepted based on whether they alter the resulting gene expression pattern to more closely resemble the target (see Materials and Methods). More specifically, we define the fitness of a mutation as the inverse of the normalized-Root Mean Squared Error (RMSE) relative to the target. Individual mutations that result in a better match to the target expression pattern (*i.e.* have lower normalized-RMSE values) are accepted, with some probability of accepting slightly deleterious mutations to allow for efficient exploration of the potentially rugged fitness landscape.

**Fig 1.**
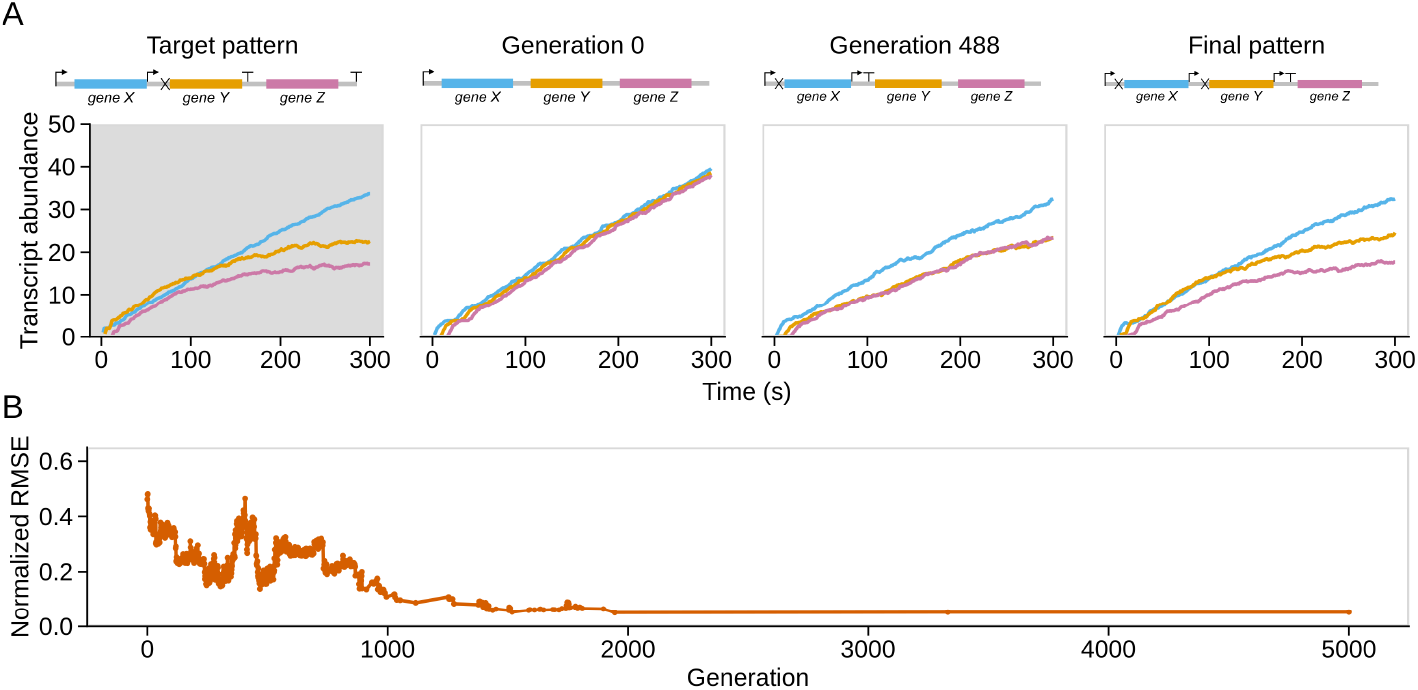
Evolutionary simulation of a positive control gene expression pattern. A: The target pattern (first panel) is a gene expression time course generated by an arbitrarily chosen genome architecture simulated in Pinetree. (Line colors correspond to the genes labeled in the genome architecture). The remaining panels illustrate the evolutionary process: the starting genome architecture and corresponding gene expression time-course (generation 0), an example from the middle of the evolutionary simulation (generation 488), and the final evolved architecture (final pattern). B: The normalized-RMSE metric declines with increasing numbers of generations as the genome architecture evolves to reproduce expression patterns that are increasingly similar to the target pattern.

As shown in Fig. 1A (right-most panel), the best gene expression time-course that evolved over a 5,000 generation evolutionary simulation qualitatively matches the target. To quantitatively make this determination, we found—through visualization of numerous simulations—that normalized-RMSE values ≤ 0.1 generally aligned with our qualitative determination about the success of any given solution. We henceforth consider an evolutionary simulation successful if the final normalized-RMSE falls below this value, which occurs in this case (Fig. 1B). Thus, this positive control shows that our approach is capable of evolving genome architectures that match complex patterns from simple starting points.

### Evolving genomes to match a range of gene expression patterns

After successfully conducting an evolutionary simulation with a positive control, we applied the same method to a range of gene expression patterns that have no *a priori* known genetic solution. We began with a simple pattern where all three genes linearly increase in abundance over time but do so at different rates (Fig. 2A, left two panels). A rational solution to this pattern might be to have a strong upstream promoter with weak terminators between each successive gene. An alternative solution would be to start with a weak upstream promoter followed by subsequent promoters between each gene so that each successive gene will be expressed at a higher level. While in the first case that we outlined the *first* gene in the genome would be expressed at the highest rate, in this latter scenario the *last* gene would be expressed at the highest rate.

**Fig 2.**
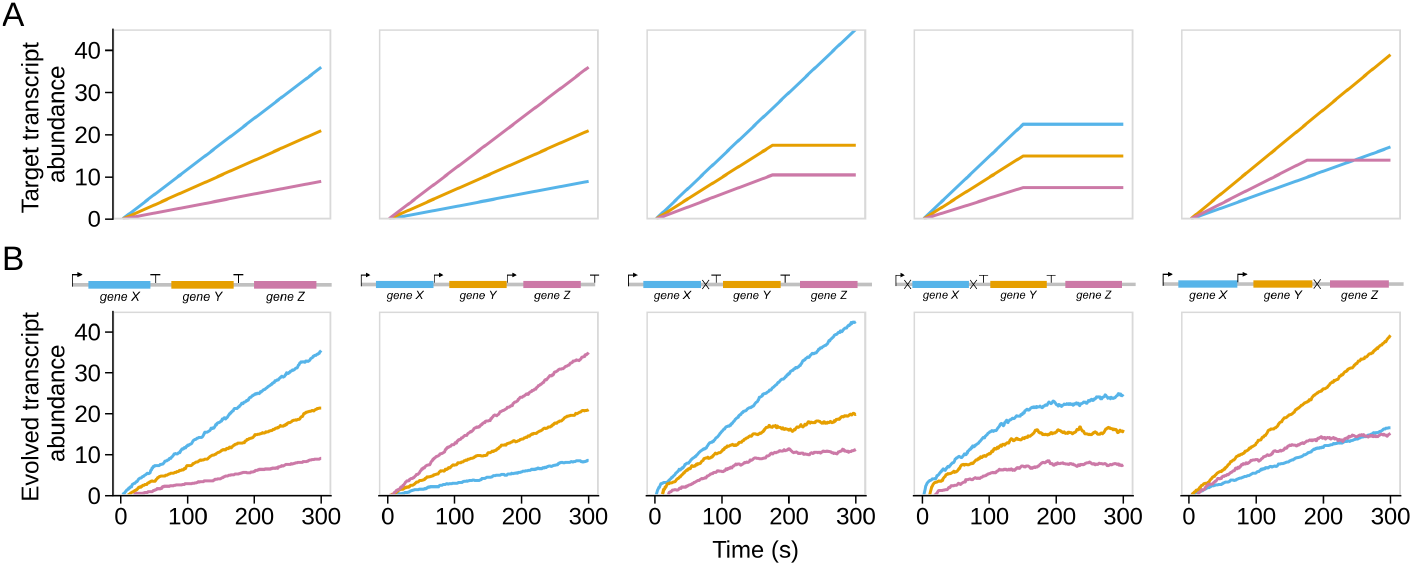
Examples of successfully evolved gene expression patterns. A: Each panel contains a target gene expression time-course pattern. Collectively, these patterns span a diverse range of possibilities. B: The corresponding representative gene expression patterns from successful simulations for each respective target. Shown above each time-course of gene expression is the evolved genome architecture that produced it.

As we expected, the left two panels of Fig. 2B show that we were able to successfully evolve this linearly increasing pattern (with two different gene arrangements) and that the evolved genome architectures match our rational expectation. The final three examples presented in Fig. 2 further show successful simulations (the lowest normalized-RMSE values found across 50 independent replicates) for more complex patterns that are difficult to rationalize solutions to. While this demonstrates that genomes in our system can evolve to match several distinct patterns with qualitatively successful solutions, we wanted to further increase the complexity of our targets and evaluate replicate simulations more quantitatively.

We conducted a total of 300 independent evolutionary simulations for each of 10 general patterns (Fig. 3A), split into 50 simulations for each of the 6 possible gene arrangements per pattern (S3 Fig, see Materials and Methods). For the particular gene arrangement that yielded the best results for each pattern, Fig. 3B shows the distribution of normalized-RMSE values from the 50 replicate simulations (S4 Fig shows corresponding gene expression time-courses for the best replicate). Each pattern had at least two successful replicate simulations (normalized-RMSE ≤ 0.1), but there was nevertheless a clear difference between how easily solutions to particular patterns were able to evolve—compare and contrast boxplots for patterns #5 and #6 in Fig 3B. For pattern #5, the box lies entirely below the red line, indicating that over 75% of simulations were determined to be successful according to our cutoff criterion. By contrast for pattern #6, the box lies entirely above the red line, indicating that over 75% of the simulations were not determined to be successful. This result suggests certain gene expression time-course patterns may have a more restrictive set of genome architectures capable of producing them, but we also note that the evolutionary parameters could perhaps be altered to better optimize solutions to particular patterns.

**Fig 3.**
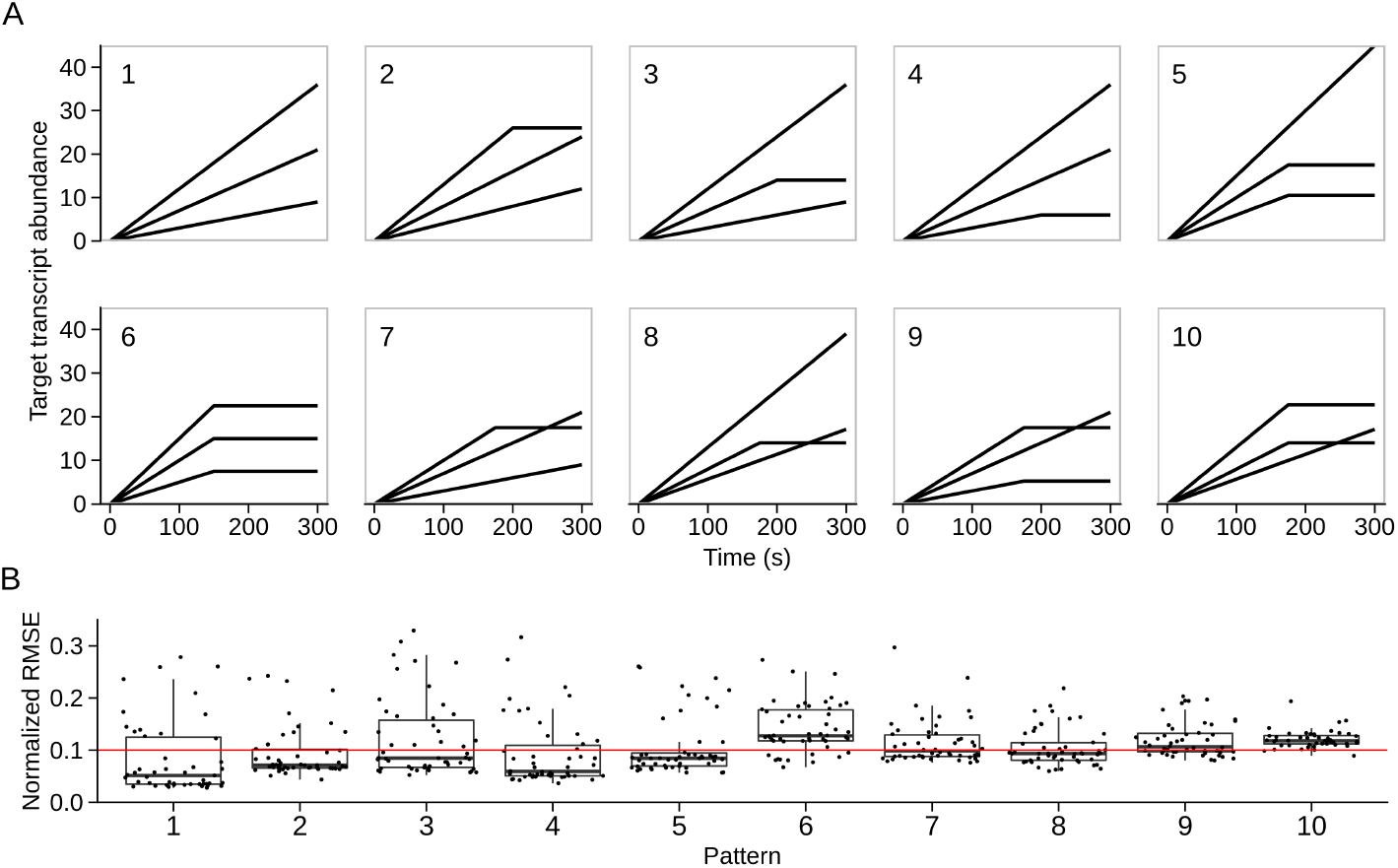
Quantitative assessment of pattern achievability. A: The 10 target gene expression time-course patterns that we simulated are displayed as black lines representing general patterns (each of which has 6 possible gene arrangements). B: For 50 replicate simulations of each pattern, shown are the lowest achieved normalized-RMSE values. All patterns had at least two successful independent evolutionary simulations, as determined by the red line highlighting a normalized-RMSE value of 0.1.

### Disparate genome architectures can recapitulate the same target expression pattern

Beginning with our positive control simulations (Fig. 1), we saw evidence that different genome architectures could produce comparable time-course patterns of gene expression. In this case, while the evolved genome produced a nearly identical gene expression pattern to the target, the evolved genome differed from the input genome that was used to produce the target expression data. To investigate this trend more broadly and for different gene expression patterns, we interrogated the diversity of the successfully evolved genome architectures for each pattern. Overall, we found that there was no single genome architecture that completely dominated results for any of the target patterns. As illustrated in Fig. 4A, even the simplest pattern that we assessed had numerous distinct (albeit qualitatively similar) genome architectures that evolved, each of which were capable of producing gene expression time-courses that successfully matched the target. However, some patterns were more heterogeneous than others in terms of the number of possible genome architectures found (Fig. 4B).

**Fig 4.**
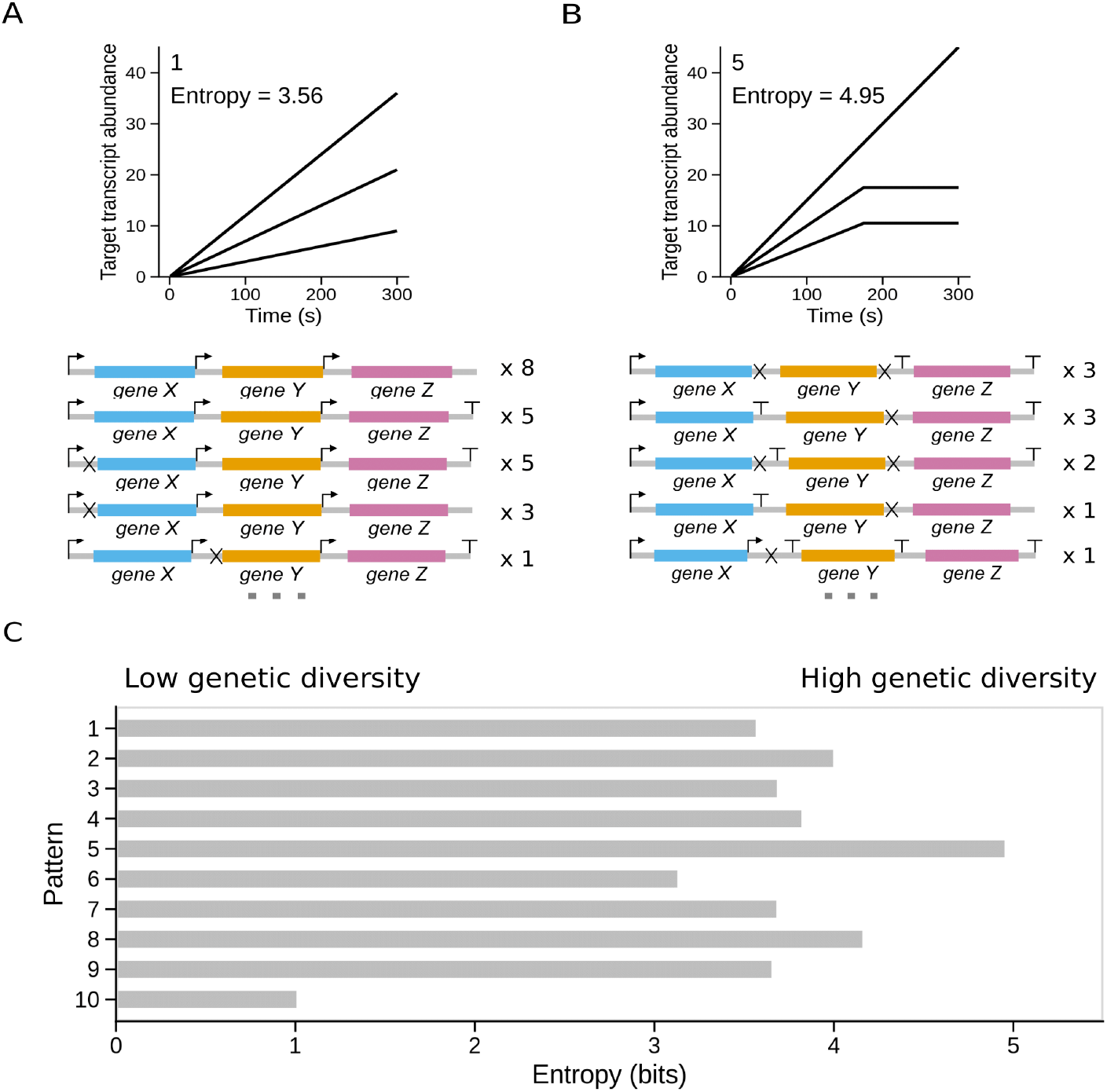
Diversity of successfully evolved genome architectures. A: A sample of the successfully evolved genome architectures for pattern #1 showing that multiple potential architectures can reproduce the target expression pattern. B: Similar to panel (A), a sample of the successfully evolved architectures for pattern #5. C: Entropy values for the set of successfully evolved genome architectures for each target pattern. Lower values indicate that successful replicate simulations converged upon one or a small number of genome architectures. More specifically, an entropy value of 1 corresponds to their being effectively 2 (2^1^) distinct architectures for a given pattern from amongst the replicates, while an entropy value of 5 corresponds to their being effectively 32 (2^5^) distinct architectures.

We quantified the diversity of evolved genome architectures for successful replicate simulations for each pattern using information entropy (as in Fig. 3 we analyzed data from only the best of the 6 possible gene arrangements for each pattern). Lower entropy values imply that only a single or a small number of distinct genome architectures evolved for a particular pattern. By contrast, high entropy values imply that the genome architecture solutions for a particular pattern were largely distinct from one another. Fig. 4C illustrates the entropy scores for each of the 10 patterns showing that pattern 5 had the most diverse set of genome solutions whereas pattern 10 (which had only two successful simulations) was the most constrained. Taken together, these results show that particular patterns may be more flexible than others in terms of the number of possible solutions but the cause of this variability is unknown at this stage and a possible avenue for future research.

### Evolutionary simulation of a ten-gene model

Although the three-gene model is ideal for studying the basic principles behind genetic regulation, the size of this genome is unrealistically small compared to most phage genomes. Thus, we wanted to test whether our simulation approach was capable of evolving larger genomes containing up to 10 genes (on par with the smallest known phage genomes). As shown in Fig. 5, we achieved a qualitatively suitable result when attempting to simulate a simple linear pattern with variable rates of increase for each gene within the genome. However, we note that increasing the number of genes within the genome also increases the run-time of the simulation substantially since more generations are required to adequately explore the fitness landscape given that there are many more possible mutations (*i.e.* regions between genes that can potentially contain regulatory elements). This result, nevertheless, shows that our approach is generalizable and can, in principle, be extended to any number of genes with complex, predefined target patterns barring computational limitations.

**Fig 5.**
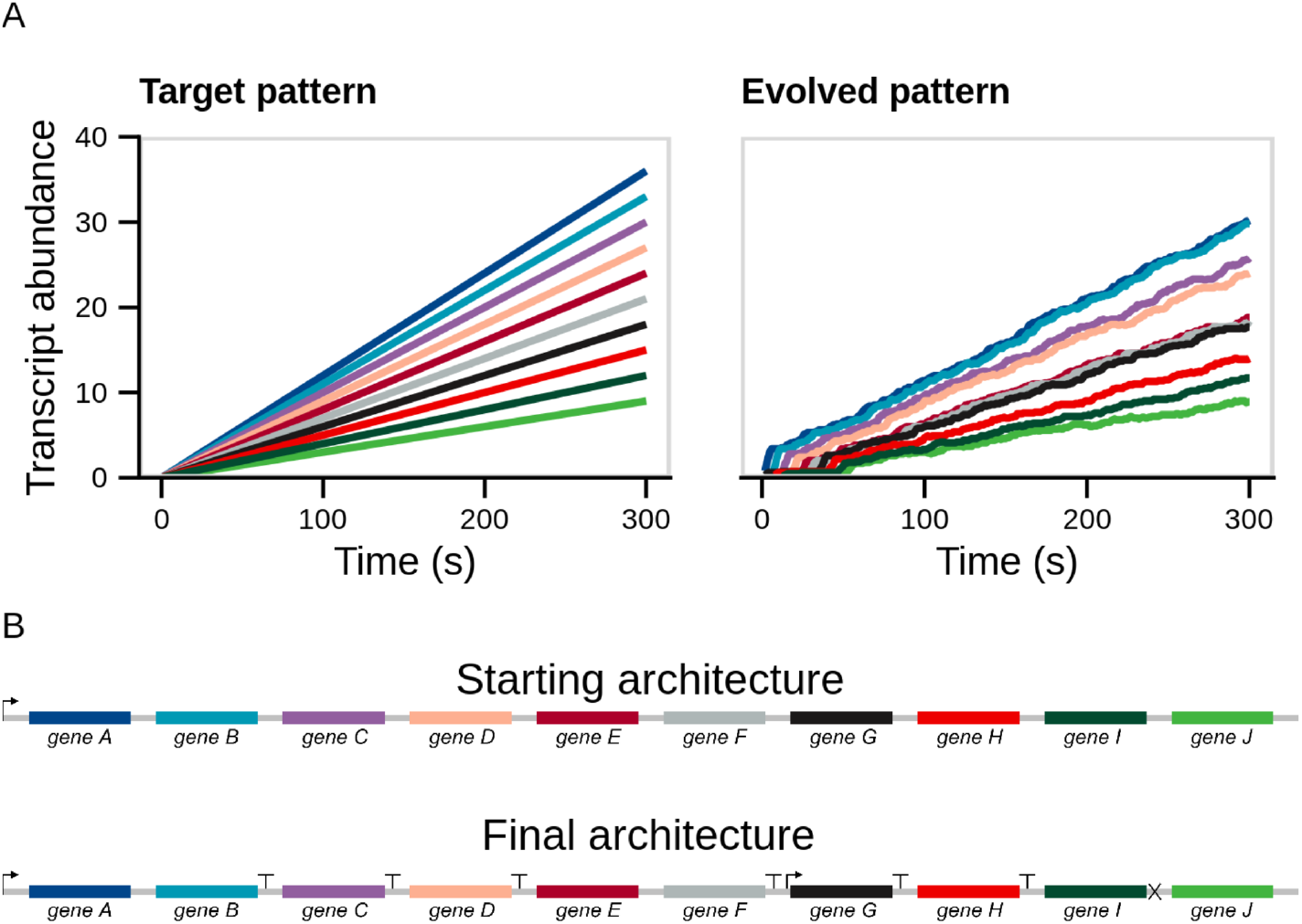
Evolutionary simulation of a ten-gene model. A: Shown are the target (left) and evolved (right) patterns. The evolved pattern is the best of 5 independent replicates, each of which was simulated for 100,000 generations. B: The starting (top) and best fitting evolved (bottom) genome architectures.

## Discussion

Genomes of all species employ a myriad of regulatory strategies to ensure that individual genes are expressed at particular levels that vary over time. Here, we have used computational modeling to demonstrate that bacteriophages can generate a range of complex and time-varying expression patterns without the need for specific regulatory molecules or complex genetic circuitry. By varying only the strength of promoters, terminators, and RNase binding sites, our *in silico* evolution platform was able to generate genomes capable of matching numerous and distinct gene expression time-course patterns. These findings show that networks of interacting promoters and transcription factors are not explicitly necessary to achieve certain gene expression designs, and provide a proof-of-principle for the *de novo* design and engineering of phage genomes using molecular-level evolutionary simulation.

Our study is motivated by the observation that naturally evolved phages appear to regulate their gene expression via the mechanisms discussed here. Phage genomes are typically small and highly gene dense, which is likely to limit the overall complexity of their native gene expression programs [31–34]. Even with relatively small genomes, numerous studies have shown that phages are capable of producing complex gene expression dynamics over the course of an infection cycle. In phage T7, for instance, genes that are expressed early in the infection cycle are typically involved in shutting down synthesis of host-cell macromolecules, whereas the genes that are expressed later have roles in viral assembly and lysis [38–40]. While the T7 genome encodes its own polymerase that is critical for the delayed expression of these late genes, phage T7 gene expression dynamics are partially accomplished through the use of variable strength promoters, transcriptional terminators, and RNase cleavage sites [27, 28, 35, 41–43]. Incorporating knowledge of variable strength regulatory elements has been critical for generating accurate computational models of phage T7 infection that are increasingly able to recapitulate empirically measured gene expression abundances [29, 35–37]. The findings presented here suggest that this level of regulatory control is likely to apply more broadly to phages in general.

Our findings here highlight some of the strengths and the limitations of regulatory approaches built solely off varying the strength of transcription, termination, and transcript degradation. In addition to accumulating at different rates over time via variable strength transcription, our system shows that gene products can also plateau at different levels by tuning degradation rates. In the most complex cases that we investigated, a combination of variable production and degradation rates is sufficient to produce expression dynamics whereby the relative ordering of genes by abundance can switch over time (Fig. 3, patterns 7–10). However, the regulatory elements that we assessed do not provide a means to shut off the production of certain genes, which would enable decreases in *absolute* transcript abundances at particular times. Similarly, gene expression delays within this system are limited to short time scales when the phage genome is still being injected; there is no apparent mechanism for inducing expression at a particular time without considering more complex genetic regulation—such as the early production of a phage-specific polymerase that controls later genes as in phage T7.

Despite these constraints, our approach may provide important insight into the overall organization and regulatory principles governing phage genomes. For instance, RNase cleavage sites and promoter sequences frequently co-occur in phage T7 [28]. In our simulations, we observed numerous instances of this co-occurrence, indicating that this motif may provide a general means for decoupling the expression of proximal genes. However, the generality and significance of this finding is difficult to assess based on the limited set of patterns that we investigated. Future work, particularly focused on larger genomes with more complex expression dynamics, might uncover interesting design principles that have heretofore gone unnoticed.

While our results provide insight into the limits and capabilities of phage regulatory strategies, our approach may also prove useful for synthetic biology applications in phages [44, 45]. Synthetic genetic circuits are typically designed for free-living species where they must operate in variable cellular contexts over comparatively long time-scales [46, 47]. By contrast, phage infections are typically brief, with relevant time-scales on the order of tens of minutes [48, 49]. We performed our simulations with realistic cellular parameters over an initial 5 minute infection period. Producing more complicated dynamics from much larger genomes over longer timescales could, in principle, be accomplished using minimal genetic circuitry to divide the genome into early and late gene sets; within each of these two sets our results show that it is possible to encode diverse dynamics without any further regulatory mechanisms. Indeed, the phage T7 genome is partitioned in precisely this manner, having distinct classes of early, middle, and late expressed genes that perform distinct functions [38–40].

In principle, the approach that we presented here could be adapted to explicitly design phage genomes with specified gene expression dynamics but doing so presents several challenges. While Pinetree relies on genome-level information, it does not make use of *sequence*-level information. Rather, elements such as promoters are encoded by their location and strength—not by an explicit sequence. Translating a particular evolutionary design into an actual genome sequence would thus require using standardized parts with known element strengths; these strengths could be calibrated against the values encoded in Pinetree to aid in genome sequence design [50]. In the case of RNase sites, this may be particularly challenging because while degradation rates are known to vary considerably across genes, the overall rules concerning RNase binding and cleavage are less well understood than the actions of promoters and terminators. However, this outlook is beginning to change [51–53].

We envision that the results presented here can be extended in a number of different directions in future research. First, we only considered non-overlapping gene arrangements in our study but many phage genomes are highly compact and successive genes frequently overlap; this feature might impact gene expression in ways that are difficult to predict but could be assessed using our approach [23, 31]. Second, an obvious way to create more complex gene expression dynamics would be to encode a regulatory protein on the phage genome rather than the arbitrary proteins that we have thus far investigated. Extensions of our framework to consider more genes would likely benefit from such an approach, which could allow for expression delays or decreases in the abundance of particular transcripts over time. Third, we previously noted that gene order can play an important role in achieving particular targets. With only three genes we were able to enumerate all possible gene arrangements, but this approach is intractable for larger genomes and will require an additional mutational step that swaps the placement of genes for larger applications. Finally, we focused our attention on transcript abundances but differences in translation initiation and elongation rates offer further ways to tune gene expression at the level of protein products to produce more complex dynamics [7, 8, 54–57].

An extraordinary number of phages have been discovered in recent years, but only a small number of these have been interrogated experimentally [58–62]. Computational approaches can help to better characterize the general constraints and principles that affect genome organization and function, and can additionally provide a way to engineer phages based on predetermined design constraints. The design and engineering of whole phage genomes using principles and approaches that we developed here may have a number of future medical and biotechnological applications, including combating antibiotic resistant bacteria [63].

## Supporting information

Combined Supplementary Information

## Supporting information

**S1 Fig. Influence of *β* parameter on the strength of selection in our evolutionary simulations.** A: We fixed effective population size at 1000 and ran 5,000 generation simulations with the *β* value varying from 0.001 to 2.000. Each plots shows representatives from 10 simulations of pattern number 1. B: We summarized panel (A) by plotting the fraction of accepted mutations that were deleterious for each *β* value. C: We settled on an approach that varies *β* over the course of each simulation (simulated annealing), with this plot showing the resulting piecewise function: *β* remains constant for the first 10% of generations, linearly increases until 90% of the generations have passed, and then linearly increases at a different rate for the final 10% of generations.

**S2 Fig. Analyzing the effective range of individual regulatory element strengths.** A: The genetic architectures above the plot display the element’s placement on the genome when analyzing their strengths. The line plot demonstrates a promoter’s effect on gene expression at differing strengths by comparing the ratio of Gene Y over Gene X (normalized transcript abundances). The grey dashed line denotes the strength at which we decided to insert a promoter in our evolutionary simulations when an ‘add’ mutation is initially proposed. B: Similar to panel (A), considering transcriptional terminator strengths. C: Similar to panel (A), analyzing RNase cleavage sites.

**S3 Fig. Example of possible gene arrangements used in the simulations.** A: The colors in the genome are used as a legend to denote each gene: blue representing gene X, orange for gene Y, and purple for gene Z. B: The six possible gene arrangements are shown for a general pattern. The need for simulating 6 gene arrangements per general pattern arises because genomic ordering of the elements may be important and our simulation does not currently include a mutational step to swap the identity of individual elements on the genome (as might happen when recombination occurs).

**S4 Fig. Gene expression time-courses from the best genome architectures found for each target pattern.** The y-axis contains each of the general patterns shown in Fig. 3 and the x-axis shows the 6 possible gene arrangements for each pattern. Each of the depicted patterns has a normalized-RMSE value below 0.1, deeming them as successful according to our threshold.

## Acknowledgments

This work was supported by National Institutes of Health grants R01 GM088344 to C.O.W. and F32 GM130113 to A.J.H. C.O.W. also received support from the Jane and Roland Blumberg Centennial Professorship in Molecular Evolution and the Dwight W. and Blanche Faye Reeder Centennial Fellowship in Systematic and Evolutionary Biology at The University of Texas at Austin. S.B.S received support from a TIDES research fellowship. The Texas Advanced Computing Center (TACC) at The University of Texas at Austin provided high-performance computing resources.

## Notes

### Competing Interest Statement

The authors have declared no competing interest.

