## Supplementary material for "Generating complex patterns of gene expression without regulatory circuits": Combined Supplementary Information

### Supplementary Information for: Generating complex patterns of gene expression without regulatory circuits

March 22, 2021

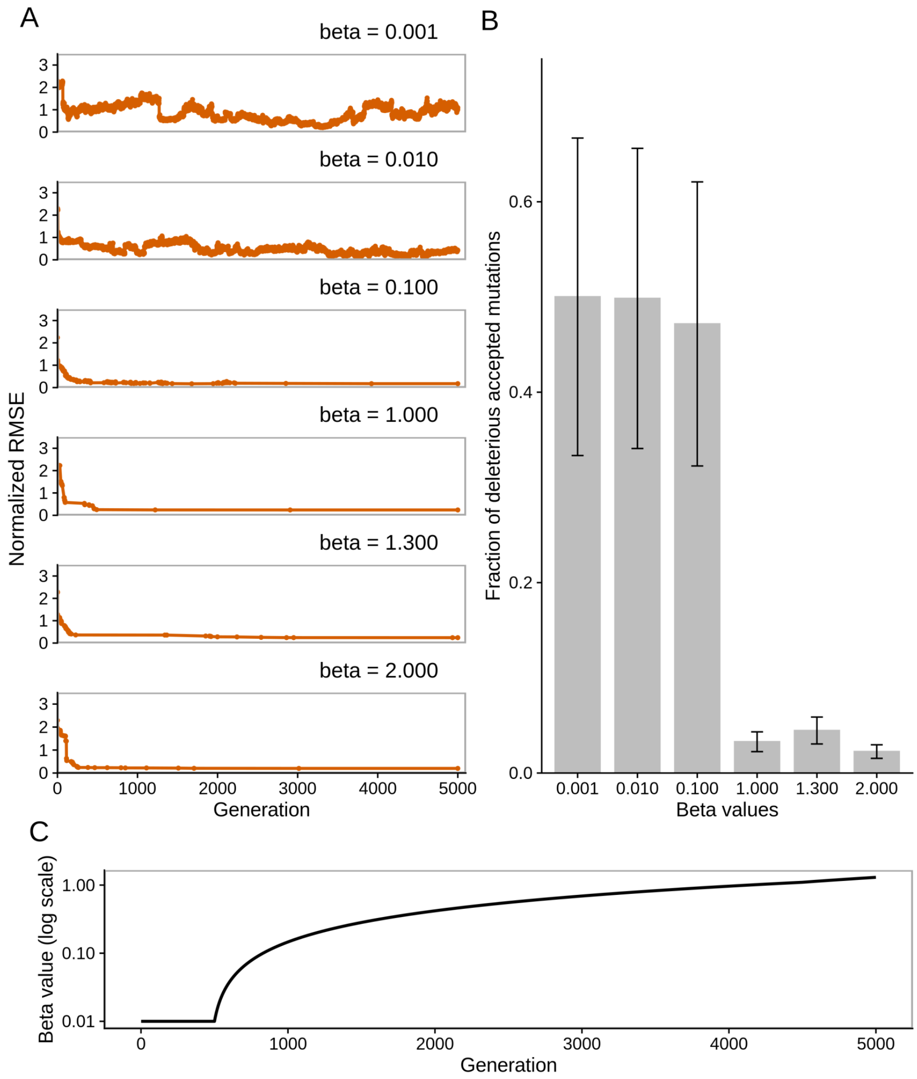


Figure S1: **Influence of $\beta$ parameter on the strength of selection in our evolutionary simulations.** A: We fixed effective population size at 1000 and ran 5,000 generation simulations with the $\beta$ value varying from 0.001 to 2.000. Each plots shows representatives from 10 simulations of pattern number 1. B: We summarized panel (A) by plotting the fraction of accepted mutations that were deleterious for each $\beta$ value. C: We settled on an approach that varies $\beta$ over the course of each simulation (simulated annealing), with this plot showing the resulting piecewise function: $\beta$ remains constant for the first 10% of generations, linearly increases until 90% of the generations have passed, and then linearly increases at a different rate for the final 10% of generations.

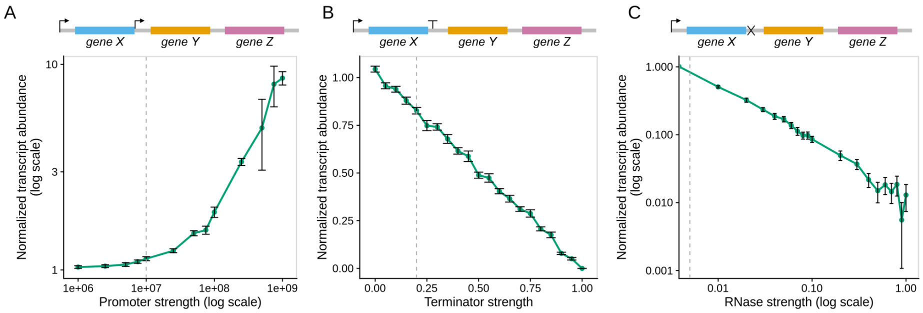


Figure S2: **Analyzing the effective range of individual regulatory element strengths.** A: The genetic architectures above the plot display the element's placement on the genome when analyzing their strengths. The line plot demonstrates a promoter's effect on gene expression at differing strengths by comparing the ratio of Gene Y over Gene X (normalized transcript abundances). The grey dashed line denotes the strength at which we decided to insert a promoter in our evolutionary simulations when an 'add' mutation is initially proposed. B: Similar to panel (A), considering transcriptional terminator strengths. C: Similar to panel (A), analyzing RNase cleavage sites.

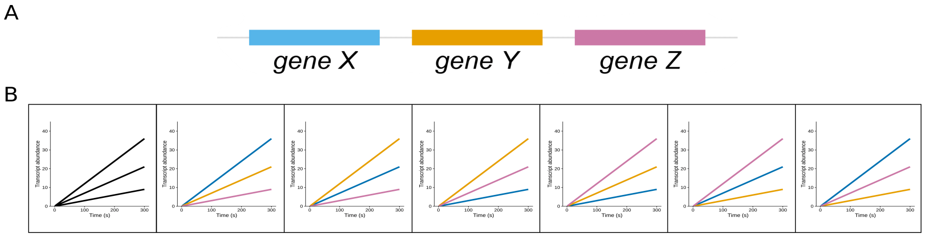


Figure S3: **Example of possible gene arrangements used in the simulations.** A: The colors in the genome are used as a legend to denote each gene: blue representing gene X, orange for gene Y, and purple for gene Z. B: The six possible gene arrangements are shown for a general pattern. The need for simulating 6 gene arrangements per general pattern arises because genomic ordering of the elements may be important and our simulation does not currently include a mutational step to swap the identity of individual elements on the genome (as might happen when recombination occurs).

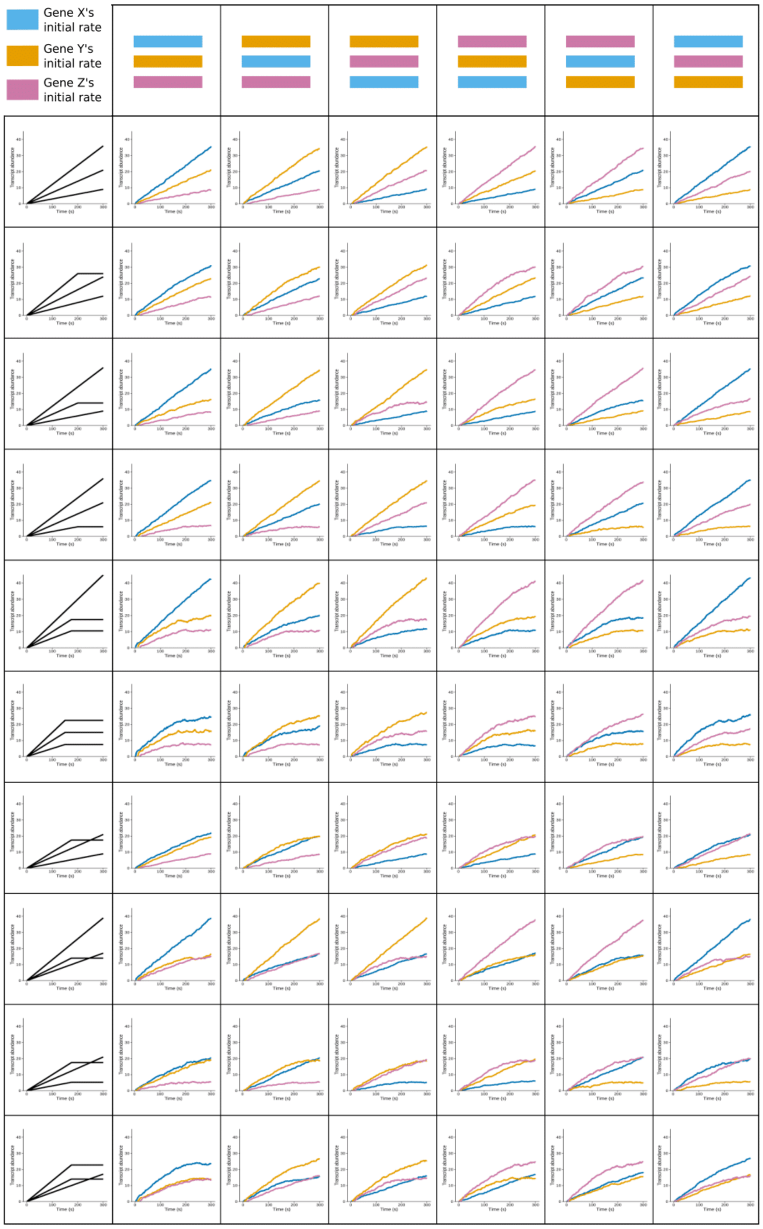


Figure S4: **Gene expression time-courses from the best genome architectures found for each target pattern.** The y-axis contains each of the general patterns shown in Fig. 3 and the x-axis shows the 6 possible gene arrangements for each pattern. Each of the depicted patterns has a normalized-RMSE value below 0.1, deeming them as successful according to our threshold.
